# Merfin: improved variant filtering and polishing via k-mer validation

**DOI:** 10.1101/2021.07.16.452324

**Authors:** Giulio Formenti, Arang Rhie, Brian P. Walenz, Françoise Thibaud-Nissen, Kishwar Shafin, Sergey Koren, Eugene W. Myers, Erich D. Jarvis, Adam M. Phillippy

**Affiliations:** The Rockefeller University and Howard Hughes Medical Institute; Genome Informatics Section, Computational and Statistical Genomics Branch, NHGRI, NIH, Bethesda, MD, USA; NCBI, NLM, NIH, Bethesda, MD, USA; UC Santa Cruz Genomics Institute, Santa Cruz, CA, USA; Max Planck Institute of Molecular Cell Biology and Genetics, Dresden, Germany

## Abstract

Read mapping and variant calling approaches have been widely used for accurate genotyping and improving consensus quality assembled from noisy long reads. Variant calling accuracy relies heavily on the read quality, the precision of the read mapping algorithm and variant caller, and the criteria adopted to filter the calls. However, it is impossible to define a single set of optimal parameters, as they vary depending on the quality of the read set, the variant caller of choice, and the quality of the unpolished assembly. To overcome this issue, we have devised a new tool called Merfin (*k*-**mer** based **fin**ishing tool), a k-mer based variant filtering algorithm for improved genotyping and polishing. Merfin evaluates the accuracy of a call based on expected k-mer multiplicity in the reads, independently of the quality of the read alignment and variant caller’s internal score. Moreover, we introduce novel assembly quality and completeness metrics that account for the expected genomic copy numbers. Merfin significantly increased the precision of a variant call and reduced frameshift errors when applied to PacBio HiFi, PacBio CLR, or Nanopore long read based assemblies. We demonstrate the utility while polishing the first complete human genome, a fully phased human genome, and non-human high-quality genomes.

## Introduction

Accurate variant calling has been a challenge in medical genomics, especially to achieve both high recall and precision in hard to measure regions^1^. The advent of Next Generation Sequencing (NGS) and long-read sequencing technologies streamlined variant calling algorithms^2^, which typically include: 1) aligning all reads to a reference genome; 2) performing variant calling from the alignment; and 3) filtering to remove false positives. The final outcome of variant calling relies heavily on the precision of this multistep procedure, which depends on: 1) the quality of the read set; 2) the precision of the read mapping algorithm; and 3) the precision of the variant caller in generating reliable calls^3^. To remove false positives, variant calls are often hard filtered using heuristics such as requiring a minimum coverage support, genotype quality, or other internal quality scores^2^. However, no consensus on the optimal parameters has been established, and the best parameters vary depending on the sequencing technology used. Therefore, the accuracy of a variant often corresponds to the theoretical limit of the algorithms and the parameters employed, and not the theoretical limit given the quality of the supporting raw data.

In parallel, new sequencing technologies greatly expanded our genome assembly toolkit. While the short read assemblies stumbled resolving repetitive regions^4^, long reads have considerably improved the contiguity of genome assemblies^5^. However, reduced consensus accuracy has been progressively acknowledged due to the lower base calling accuracy in long reads until the more recent PacBio Hi-Fidelity (HiFi) reads became available^6^. Still, lower base call accuracy remains even in HiFi reads for simple repeat sequences, particularly homopolymers^7,8^. Reduced consensus accuracy has detrimental impacts on many downstream analyses, such as gene annotation, which requires an accurate consensus to predict the correct coding sequence^7^. To mitigate this issue, “polishing” tools have been developed, such as Pilon, Arrow, Racon and Medaka^9–11^, while established variant calling tools such as GATK, Freebayes, DeepVariant^12–14^ have been repurposed to detect errors and find candidate corrections. Unlike re-sequencing based methods, the assembly from the same genome is used as a reference for polishing, and thus all homozygous variants suggest corrections to be made. Once corrections are collected, the consensus can be updated using tools such as Bcftools^15^. The process is usually repeated with different read sets (e.g. long and short reads), until the accuracy of the consensus reaches a set standard.

Consensus accuracy (hereby noted as QV for simplicity) has been historically measured from the variant calling process as described above, however, bearing biases caused from mapping or variant calling. In our previous work, we presented Merqury^16^, an alignment-free approach to estimate base-level QV using *k*-mers (genomic substrings of length k). In Merqury, k-mers found only in the assembly and not in the reads are considered as errors, disregarding the expected copy number. As a result, overly represented *k*-mers from sequence expansion (i.e. false duplications) in the assembly are considered as correct bases. Merqury also presents a completeness metric from the portion of *k*-mers found in the assembly from a given reliable *k*-mer set in reads. However, this *k*-mer completeness metric does not account for the *k*-mer multiplicity in the reads, limiting the scope to be measured in the non-repetitive k-mer space. As a result, any two assemblies with the same distinct *k*-mers will score the same completeness metric, regardless of one having higher sequence collapses or expansions.

Ideally, the sequence of an error-free and complete genome assembly is in perfect agreement with the sequence data, assuming that genomic DNA is randomly sampled and sequenced with negligible sequencing biases. Therefore, any changes introduced during polishing should improve the assembly-read agreement. This principle has been widely used to visually evaluate genome assembly copy number spectrum (e.g. copy-number spectrum analysis^16,17^), and more recently, used to detect errors and improve read alignment^18–20^. However, none of the evaluation metrics or polishing methods have fully utilized this assembly-read agreement to date.

Here, we first introduce a k-mer based filtering approach applicable on genomic variant calls, which achieved higher F1 scores compared to parameter based hard-filtering methods. Next, we propose a revised QV and completeness score that accounts for the expected sequence copy number given a *k*-mer frequency, driven by our refined *K^*^* definition^21^ for genome assembly evaluation. Our *K^*^* enables the detection of collapses and expansions, and significantly improves the QV when used to filter variants for polishing. We applied this approach to polish and evaluate the most complete HiFi-based assembly of human CHM13^22–25^, a Nanopore-based trio assembly from the Human Pangenome Reference Consortium (benchmark paper in preparation), and three non-human reference genomes from the Vertebrate Genomes Project (VGP)^5^, all resulting in significantly higher consensus accuracy and annotation quality. This approach is implemented as *Merfin* (*k*-**mer** based **fin**ishing tool) and is publicly available.

## Results

### Variant call filtering for higher precision

A reference genome (i.e. GRCh38) with its sequence replaced at all alternate variant calls can be considered as a “consensus” sequence and evaluated with k-mers. Unlike using a *de novo* assembled genome of the same individual as a reference, natural biological differences between the sequenced individual and the reference genome or the incomplete state of the reference (i.e. missing a segmental duplication) imposes challenges to reliably call variants. Nevertheless, it is possible to construct consensus paths from a variant or series of variants within *k* bps, and confirm its validity. We can score each path by the number of *k*-mers never found in the reads (error *k*-mer), and choose the best path to contain minimum error *k*-mers (**Fig. 1a**).

**Fig. 1.**
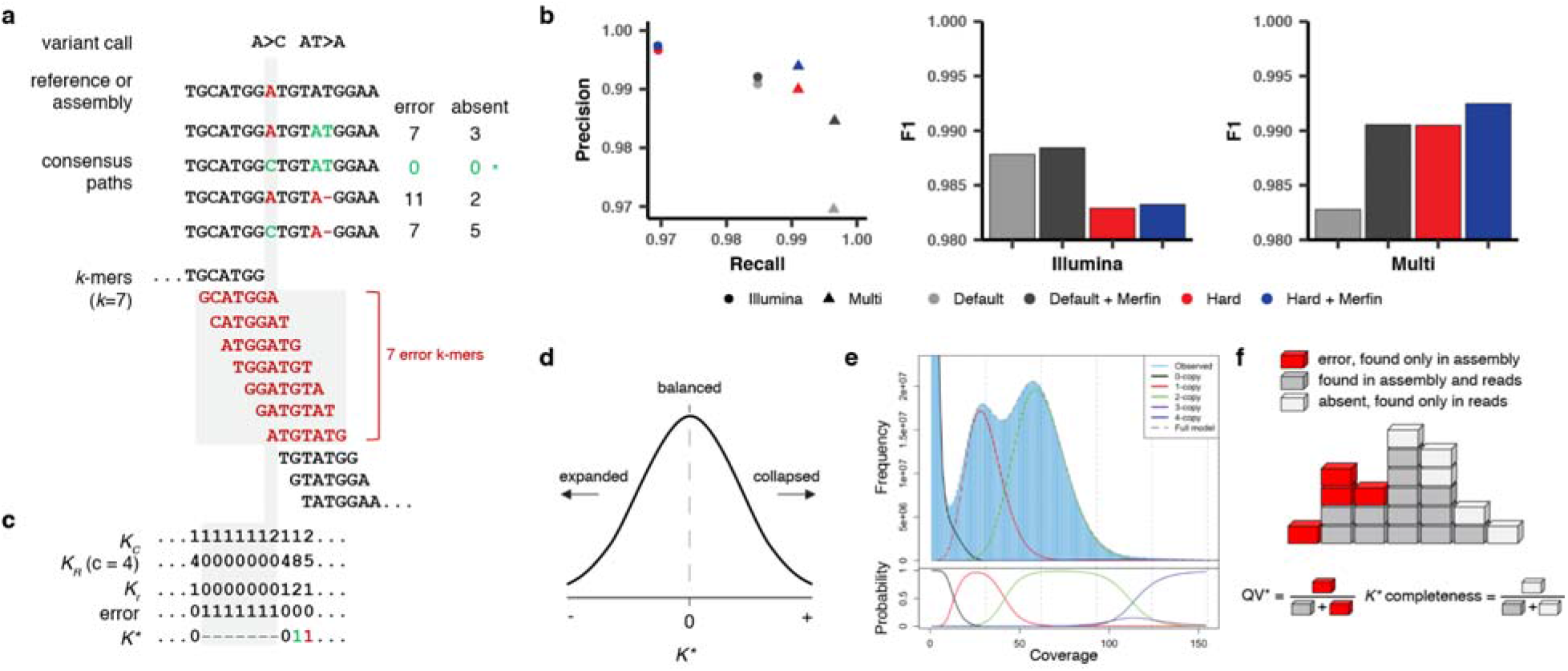
Algorithms and results used in Merfin. **a**, Two variant calls and its potential consensus paths. The bases and k-mers in red are errors not found in the reads. The path with A>C has no error k-mers and gets chosen for genotyping (*). For polishing, the average *K^*^* gets computed in addition to the missing k-mers using the predicted absent k-mers. **b**, Precision, recall, and F1 from a benchmark on HG002 genotyping. Merfin always achieves higher precision and F1 scores compared to the hard-filtered approach with almost no loss in recall. Default, no filtering; Red, hard-filtering on default. **c**, K-mer frequency found in the consensus sequence (K_C_), reads (K_R_) with averag coverage at 4 (c), expected copy number based on the corrected k-mer frequency (K_r_ = K_R_ / c), and K*. Positive an negative K* values are colored in green and red. The highlighted region (gray) shows the same k-mers and values used to compute K* as affected by the A base in the reference. If two alternatives bear the same number of missing k-mers the alternative with the K* closest to zero is chosen. **d**. K* distribution. K* values deviated from 0 indicate collapsed (+) or expanded (-) k-mers in the assembly. **e**, Genomescope 2.0 k-mer frequency histogram with theoretical k-multiplicity curves (top) and probabilities (bottom) for 0, 1, 2, 3, and 4-copy k-mers, generated using the `--fitted_hist` option. Note that the 3-copy peak is fully contained in the 2 and 4-copy peaks. **f**, Diagram for estimating QV* and completeness from k-mers. Each k-mer is a block colored by its state of presence. In the block tower, each column represents the identical k-mer with its state colored by its presence in the assembly, reads, or in both. Note the QV^*^ and K^*^ completeness is using all k-mers including their frequency.

To test the validity of this filtering approach, we benchmarked against unfiltered (default) and hard-filtered variant calls submitted to precisionFDA challenge II, HG002^1^. The variants were called from Illumina reads or from multiple platforms (Illumina, PacBio HiFi, and ONT) using GRCh38 as the reference with GATK HaplotypeCaller. Hard-filtering was performed using the variant caller’s internal scores such as PASS, QD, MQ and QUAL. When comparing precision, recall, and F1 (harmonic mean of precision and recall) on a truth set of Chr. 20^26^, Merfin always achieved higher precision with minimal loss in recall when applied on both default and hard-filtered sets (**Fig. 1b**). The hard-filtered set had a higher precision, with the price of losing more true positives, resulting in a lower F1 score when compared to the default set. On the other hand, Merfin was able to remove additional false positives on the hard-filtered set.

### Assembly evaluation

When a reference genome is constructed from the same individual, the k-mer multiplicity seen in the reads is expected to match the reference. This assumption can be used for evaluating de novo assembled genomes. In the following, we introduce our revised *K^*^*, which we used to identify possible collapsed and expanded regions in an assembly, and quantitative metrics for representing the copy-number concordance and completeness of the assembly.

#### Identifying collapsed and expanded regions

The *K^*^* metric was defined previously to detect identical collapsed repeats on each *k*-mer in the assembly^21^. The method proposed *K* = K_R_ /K_C_*, where *K_R_* is the frequency of a *k*-mer found in the reads; and *K_C_* is the frequency of a *k*-mer across the entire consensus sequence of the assembly. In regions with no collapsed repeats, *K^*^* will be equal to c, the average coverage of the sequenced reads. Here we revised the *K^*^* such that it evaluates both collapses and expansions. We propose *K^*^* = (*K_r_ - K_C_*) / min(*K_r_*, *K_C_*), where *K_r_* is the expected copy number inferred from the reads (**Fig. 1c**). For a perfect genome assembly and an unbiased read set, *K^*^* is normally distributed with mean 0, and deviations from the mean reflect natural variation in the Poisson sampling process (**Fig. 1d**). Conversely, any significant deviation from the normal distribution can be interpreted either as a bias in the assembly (i.e. an assembly error) or a bias in the read set. In particular, a positive *K^*^* implies that the assembly contains fewer copies of *k*-mers than suggested by the read set (collapsed), while negative *K^*^* implies more copies in the assembly than suggested by the read set (expanded).

The *K_r_* can be obtained with ⍰ *K_R_* / *c* ⍰, where *c* is the haploid (1-copy) peak of the k-mer distribution of the reads. Here we assume that rounding *K_r_* is sufficient to account for the standard deviation associated with the Poisson process underlying read generation. While this is true in the case of a perfectly sampled sequencing set, the validity of this generalization is challenged in the presence of sampling bias, systematic error in the reads, and variable degrees of heterozygosity which prevents the clear distinction of each ploidy peak. To account for this uncertainty and improve the accuracy of the results, we modified Genomescope2^27^ to probabilistically infer *K_r_* for each *K_R_*, using the observed *k*-mer count distribution in the read set. If supplied, Merfin will use these probabilities for *K_r_* < 4. (**Fig. 1e**).

#### QV* estimation

An average genome-wide QV accounting for excessive copy numbers (hereby defined as QV*) can be obtained using ∑ *K_C_* - *K_r_* as errors when *K_C_* > K_r_ for all positions in the assembly (**Fig. 1f**). These excessive and error *k*-mers can be generalized as ‘errors’ and Phred-scale QV, obtained as implemented in Merqury^16^ or YAK^28^.

#### Assembly completeness

The sum of *K_r_* - *K_C_* (over all positions where *K_r_* > *K_C_*) expresses absent *k*-mers that should be present in the assembly, and can be directly translated into a measure of assembly completeness as 1 - ∑ (K_r_ - K_C_) / K_r_ (**Fig. 1f**). Importantly, contrary to other measures of assembly completeness based on a subset of the *k*-mers (e.g. relying only on the occurrence of distinct *k*-mers as implemented in Merqury16), Merfin uses all k-mers, including its frequency, and computes the fraction of the expected total number of k-mers.

### Sequence polishing

The *K^*^* becomes particularly useful in polishing. The impact of each correction or combination of corrections are assessed from a given variant call set (correction candidates) by comparing the change in *K^*^*-related metrics (**Fig. 1a,c**). In addition to the error *k*-mers collected in each predicted consensus path, we compute the consequent *k*-mer frequency change, and choose the correction only when it improves the assembly-read agreement. For example, when a suggestive correction (replacing AT with A as shown in **Fig. 1a**) introduces more error *k*-mers, it should not be used for polishing. Even when no error *k*-mers are introduced, *K^*^* theoretically informs whether a path improves the assembly-read agreement in polishing. The current implementation evaluates each path independently, and thus only a local optimum is guaranteed. Moreover, variants within distance k are considered in all combinations, allowing Merfin to filter ambiguous variant calls. This approach is fully independent of the raw dataset employed. For instance, the assembly could be generated using long reads, and the calls evaluated using either short or long reads or both, taking advantage of the strengths of each sequencing platform, making accurate orthogonal validation possible, and maximizing the assembly-read agreement.

We first focus our evaluation of Merfin on the most complete assembly to date, generated from a nearly homozygous diploid CHM13hTERT (CHM13) cell line, simultaneously released by the Telomere-to-Telomere (T2T) Consortium^22^. We then provide an example of improved base calling accuracy in a Nanopore-based trio assembly from the Human Pangenome Reference Project (HPRC). We further extend the usage of Merfin to haploid and pseudo-haploid assemblies applying Merfin to three long-read assemblies (a fish, reptile, and bird) generated in the framework of the Vertebrate Genomes Project (VGP)^5^.

### Evaluating a complete human genome: T2T-CHM13

The CHM13 cell line originates from a complete hydatidiform mole (46, XX), where both haplotypes are near-identical except for a few heterozygous variants that probably have their origin in the original mole tissue or have accumulated during cell passages^29^. This cell-line was used to generate the most complete high-quality human reference to date, resolving all centromeric and telomeric repeats and all segmental duplications and satellite arrays^22,23^. Notably, the T2T-CHM13v0.9 was polished from a variety of variant calls, with an earlier version of Merfin and improved the consensus accuracy of the final assembly^25^. We further evaluated candidate assemblies to identify collapses and expansions using Merfin using *k*-mers from HiFi and Illumina reads. We found that the T2T-CHM13v1.0 assembly shows a remarkable agreement with the raw data, with only a few regions having *K^*^* significantly different from 0, coinciding with satellite repeats (**Fig. 2a**). Rather than being assembly errors, these disagreements were associated with context-dependent augmentation or depletion in HiFi and GC bias in Illumina^22,25^. Indeed, *K^*^* derived from HiFi and Illumina *k*-mers showed opposite behavior in some regions, i.e. the HSat3 of Chr. 9 (**Fig. 2b**). These effects originating from sequencing biases were observed only on the highly repetitive regions of the genome.

**Fig. 2.**
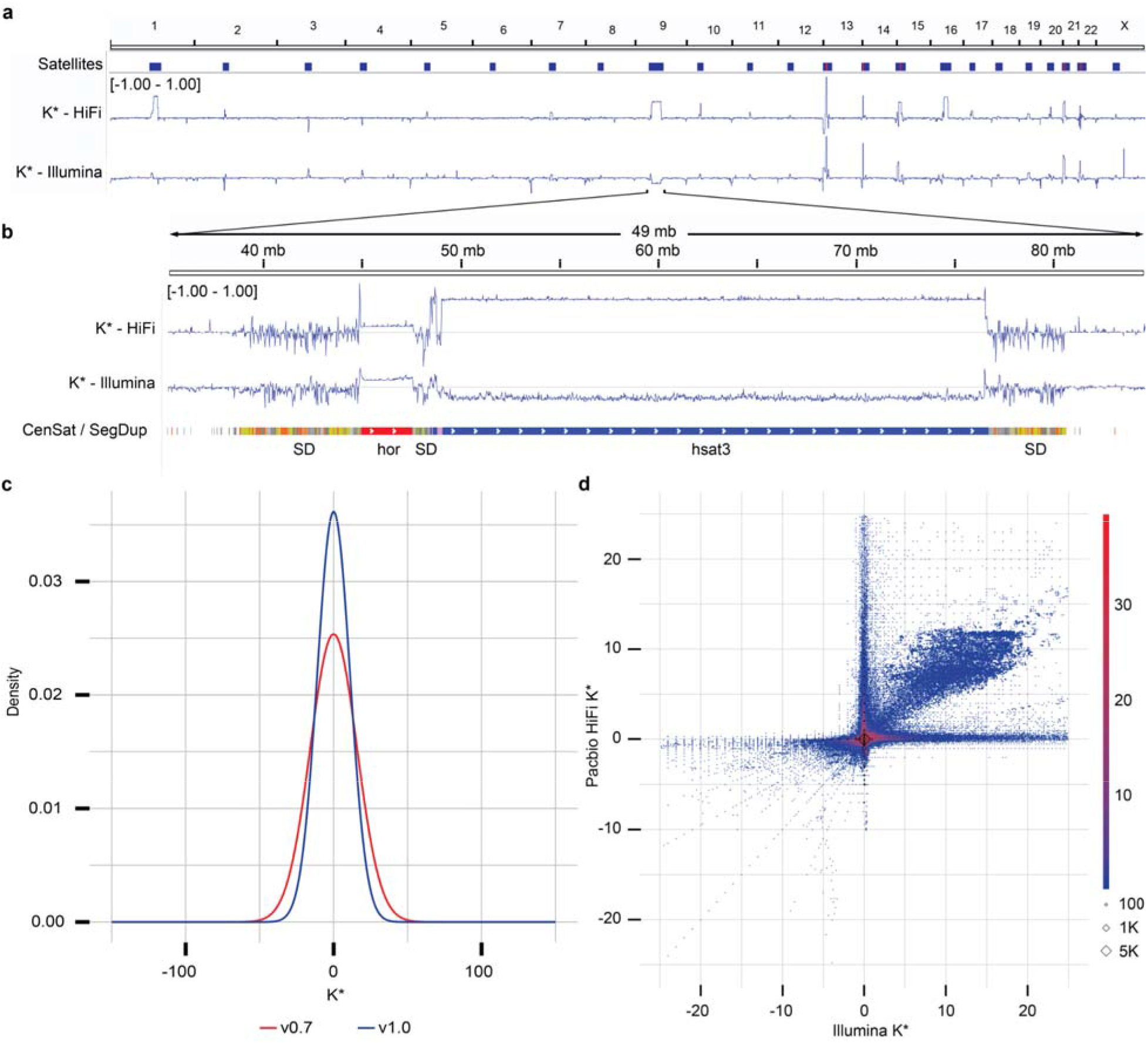
CHM13 evaluation and polishing. **a**, Genome-wide K* for the CHM13 assembly v1.0. Satellites ar associated with repeat- and technology-specific biases. Yet to be resolved rDNA arrays (red) are highlighted by positive K*. **b**, Highlight of the centromeric satellite repeats (manuscript in preparation) and segmental duplications^23^ (orange most similar, yellow less, gray least) on chromosome 9. **c**, Genome-wide density distribution of the K* using HiFi *k*-mers. When the assembly is in agreement with the raw data, the K* is normally distributed with mean 0, and the smaller the standard deviation the higher the agreement. CHM13 v1.0 shows a less dispersed distribution of the K* compared to a less complete v0.7 assembly. **d**, Genome-wide comparison of the K* computed using HiFi vs Illumina *k*-mers on the CHM13 v1.0 assembly. Agreement between the assembly and the raw reads supported by the two technologies is found around (0, 0). The upper right quadrant highlights where both HiFi v Illumina technologies suggest the presence of underrepresented *k*-mers that were mostly contributed from the un-assembled rDNAs later resolved in v1.1^22^; the lower left quadrant highlights where both technologies suggest th presence of overrepresented *k*-mers (with perfect agreement found on the diagonals). The axes correspond to regions of substantial disagreement between the two technologies. Diamonds indicate *k*-mers missing from one (x or y axis) or both (0, 0) technologies.

Compared to a less complete and less accurate preliminary assembly, T2T-CHM13v0.7^30^, T2T-CHM13v1.0 had a higher agreement of the assembly with the *k*-mers derived from HiFi (**Fig. 2c**) and Illumina (**Supplementary Fig. 1**) reads. We found a general agreement in *K^*^* between HiFi and Illumina PCR-free *k*-mers, including regions with sequencing bias common to the two technologies (**Supplementary Fig. 2**). In other cases, the direct comparison of the *K^*^* computed from the two technologies highlighted technology-specific sequencing biases (**Fig. 2a,d**). At base resolution, the *K^*^* could distinguish regions with accurate consensus from base pair errors, small and large indels, heterozygous sites, and collapsed/expanded regions (**Supplementary Figs. 3a-b**).

Both QV and QV* measured with Merqury and Merfin improved from v0.7 to v0.9 ^25^, which involved a complete reassembly of the genome using HiFi reads and patches from v0.7 at GA-rich sequence dropouts in the HiFi reads (**Supplementary Table 1**). Merqury QV improved from v0.7 to v0.9, due to the dramatic decrease in error *k*-mers, however the Merfin QV* did not change as the number of error *k*-mers is small compared to the number of overly-represented *k*-mers. Although the high amount of overly represented k-mers both in HiFi and Illumina reads may originate from sequencing biases, we argue that QV* may be a more reliable metric, because it accounts for all expected *k*-mer copy numbers, reflecting the full extent of genome representation. The QV* was also marginally influenced by the coverage through a titration experiment (**Supplementary Fig. 4**, **Supplementary Table 2**).

### Polishing a completely phased assembly: HG002

The need for polishing is particularly evident in genome assemblies generated using noisy long reads. Therefore, we tested Merfin’s variant calling filtering algorithm on a Nanopore-based assembly of human HG002 trio data generated by the HPRC using Flye^31,32^. We benchmarked Merfin on Medaka, by comparing polishing outcomes from Medaka with or without filtering with Merfin. In a trio setting, the optimal approach is to polish each parental assembly separately, by aligning the binned reads and performing variant calling^5,33^. This will prevent, or at least significantly reduce, the introduction of haplotype switches. However, our *k*-mer based evaluation of the corrections is best performed on a combined assembly so that it faithfully represents the expected copy-number of each *k*-mer given the read set. This further guards against haplotype switches since, even in case of read misbinning or mismapping, *k*-mers from the other haplotype will not be considered underrepresented.

Therefore, we first called variants separately from the binned reads used in the assembly with Medaka, and then combined the variant calls and the parental assemblies for Merfin. We conducted five different experiments using read sets that differ in coverage, version of the Guppy basecaller, and read length cut-off (**Supplementary Table 3**). Two rounds of polishing were performed in all experiments, with the second round performed on the consensus from the first round generated with the additional Merfin step. Overall, in all experiments we observed comparable improvements in base calling accuracy as measured by Merqury QV when Merfin filtering was applied (**Supplementary Table 3**). This increase reflected a dramatic positive shift in the QV distribution of individual contigs, with most low-quality contigs being rescued by Merfin, and a sharp increase in the number of contigs found without errors, leading to a final Q43.2 and Q42.8 for maternal and paternal haplotypes, respectively (**Fig. 3a)**. In the second round of polishing, the QV ceased to improve or even decreased when Merfin was not applied (**Fig. 3a**, **Supplementary Table 3**), suggesting that the best trade-off between errors corrected and introduced in the assembly was already reached in the first round. In contrast, the QV continued to increase relative to the first round with Merfin. Haplotype blocks as defined by Merqury increased in a comparable if not better way when using Merfin (**Fig. 3b**), while the haplotypes remained fully phased (**Supplementary Fig. 5**). Importantly, the results with Merfin were achieved by introducing only a fraction of the variants proposed by Medaka, making this approach more conservative than the regular polishing (**Fig. 3c**).

**Fig. 3.**
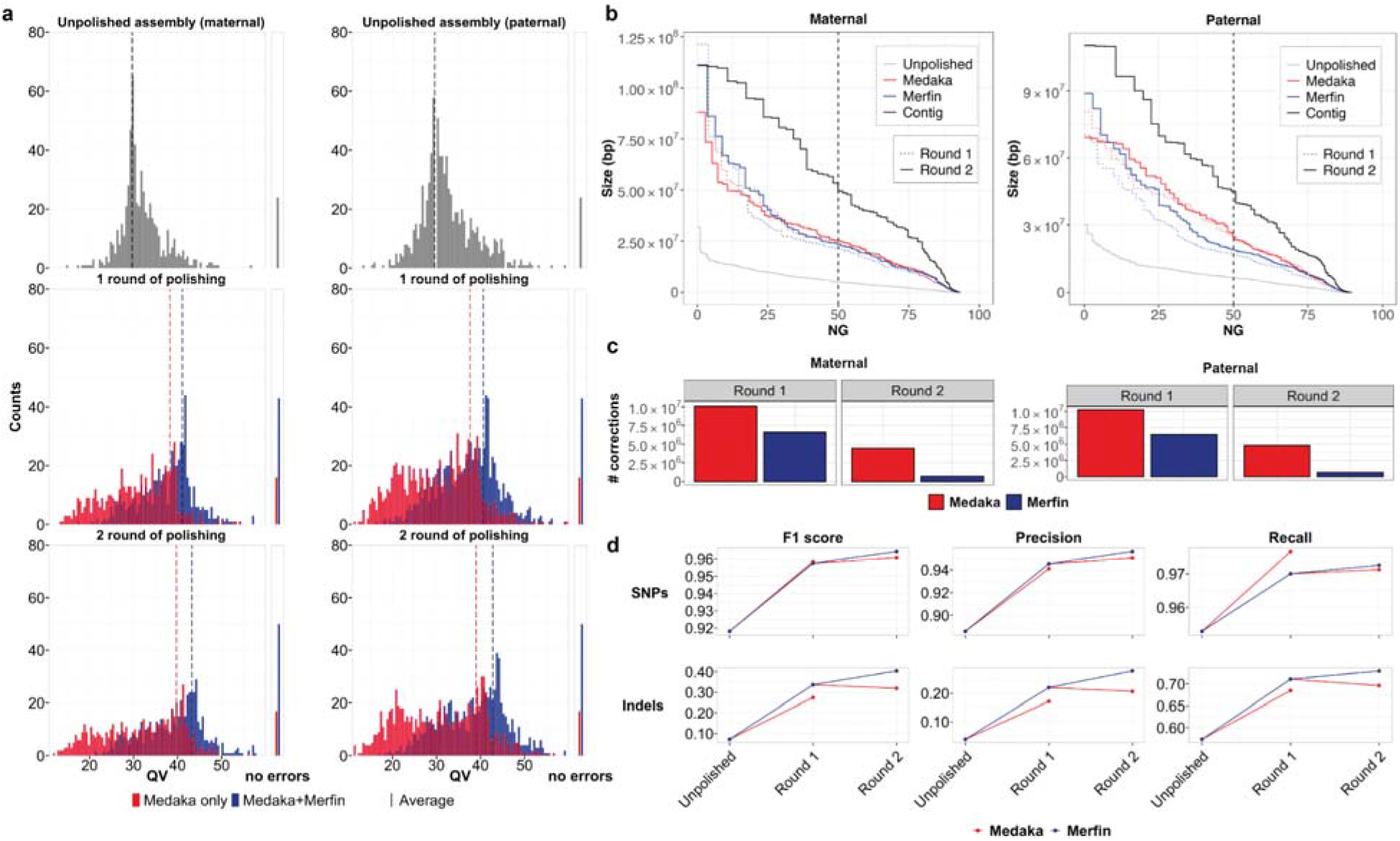
HG002 human trio polishing and evaluation. **a,** Distribution of QV scores as measured by Merqury for maternal and paternal contigs polished with Medaka only, or with variants generated by Medaka filtered with Merfin, from the experiment using latest basecaller and highest coverage. The first panel represents the unpolished contigs, the mid panel the first round of Medaka polishing and filtering, and the last panel the second round applied to the Merfin results from the previous round. The number of contigs without evidence of errors as judged by Merqury QV are reported on the right side. **b**, Size of the haplotype blocks before and after polishing with or without Merfin for both the maternal and paternal assemblies. First round of polishing is represented by the dotted lines. **c**, Number of variants generated by Medaka for polishing and remaining variants after Merfin filtering for both the maternal and paternal assemblies. **d**, Assembly-based HG002 small variant calling performance of Merfin vs regular Medaka against GIAB truth set. Variants from the assembly are derived against GRCh38 using dipcall.

We also validated the HG002 unpolished and polished assembly by aligning each haplotype assembly to GRCh38 and deriving small variants. When benchmarked against GIAB v4.2.1 truth set^26^, the results show that using Merfin we get a better F1-score, particularly when INDELs are considered (**Fig. 3d**, **Supplementary Table 4**)^26,34,35^.

### Evaluation, polishing and annotation of pseudo-haploid assemblies

We next applied Merfin to the polishing steps of the VGP assembly pipeline^5^ (**Supplementary Fig. 6**) on pseudo-haplotype assemblies from three species (flier cichlid, *Archocentrus centrarchus*, fArcCen1; Goode’s desert tortoise, *Gopherus evgoodei*, rGopEvg1; and zebra finch, *Taeniopygia guttata*, bTaeGut1). Using Pacbio continuous long reads (CLR) and 10x Genomics linked-reads for polishing, we observed general improvement in QV as measured by Merqury (**Fig. 4a**, **Supplementary Table 5**). The largest improvement was observed in the first round of Arrow polishing step using CLR. Arrow can replace low quality sequences with patch sequences generated *de novo* from the reads that align to the region, i.e., independent from the quality of the original reference. We observed low coverage, sequencing biases (i.e. homopolymer shortening), and mosaic haplotypes in the generated patches, leading to cases of lower QV in the polished assembly (e.g. **Fig. 4a**, rGopEvg1). In all cases tested, Merfin rescued the QV decrease, or improved the QV. In the subsequent polishing steps performed using Freebayes as the variant caller, the benefit of running Merfin to filter the variant set was less pronounced but still present (**Fig. 4a**, dashed lines). This was true in all cases but the zebra finch, where the default pipeline performed marginally better. However, when considering low frequency *k*-mers as errors from the probability model in Merfin, the QV as well as QV* increased in all cases (adjusted QV and QV* in **Fig. 4bc**, **Supplementary Table 5**). Merqury QV counts all *k*-mers never seen in the reads as errors, while the adjusted QV additionally counts low frequency *k*-mers based on the *k*-mer frequency spectrum as errors. The QV* further includes overrepresented *k-*mers as errors. In doing so, the QV* captures not only the base accuracy errors, but also false duplications, expressing the uncertainty associated with any particular base given the support from the raw reads.

**Fig. 4.**
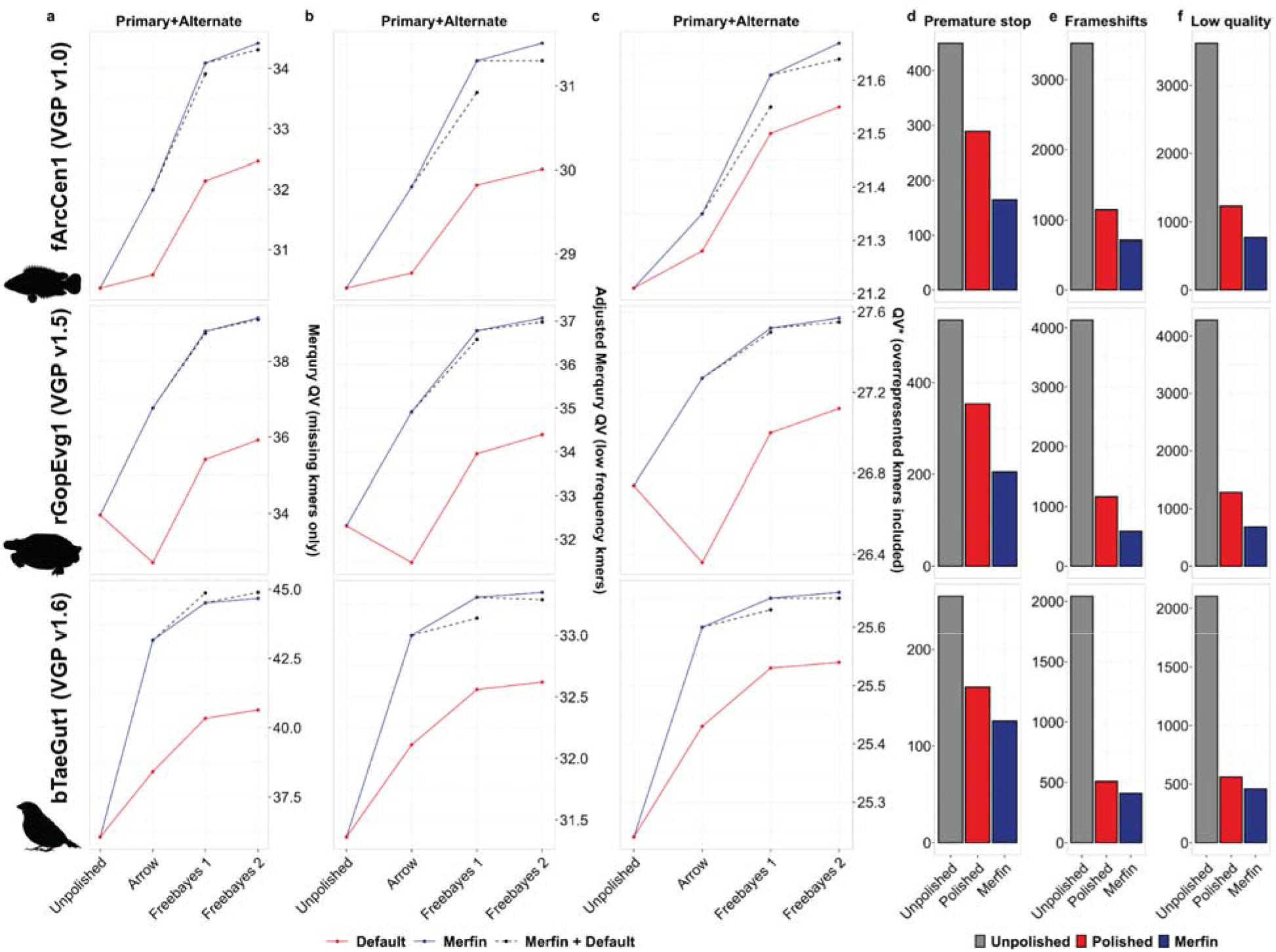
Polishing and evaluation of VGP pseudo-haploid assemblies. **a-c**, Polishing results of primary an alternate assemblies for the flier cichlid (fArcCen1), the Goode’s desert tortoise (rGopEvg1), and the zebra finc (bTaeGut1) using the VGP pipeline. Graphed are the unpolished QV values, and the Merqury QV that accounts onl for missing *k*-mers (a), the Merqury QV corrected using Merfin models for 0-copy *k*-mers (b), and QV* that als accounts for overrepresented *k*-mers (c). **d-f**, the general QV increase was reflected in the quality of the gene annotation, with consistent reduction in the number of genes affected by premature stop codons (d), frameshifts errors (e), and low quality protein-coding gene predictions (f).

Most long-read assemblers naturally generate locally phased haplotypes (e.g. Falcon-Unzip^36^), and it is therefore important that the polishing does not introduce haplotype switches. To test whether the increase in QV from Merfin was due to introducing haplotype switches, we tested a zebra finch (*Taeniopygia guttata*, bTaeGut2) pseudo-haplotype assembly for which parental sequence information is available^5^. When Merfin was applied to filter variants generated by Freebayes on the Longranger alignments of the 10x reads, we noticed an increase in the number of haplotype switches as measured with Merqury (**Supplementary Table 2**). We realized that this was due to many heterozygous variants being called by Freebayes, when individual reads were mapped to collapsed regions or preferentially to the primary assembly which had higher base accuracy^5^. Indeed, such assemblies violate the read-assembly agreement, potentially leading to heterozygous calls being preferred by Merfin. Even in almost complete pseudo-haplotype assemblies, short reads can be easily mismapped, leading to spurious heterozygous calls. To overcome this issue, we decided to remove all heterozygous variants before applying with Merfin. This substantially prevented haplotype switches (**Supplementary Fig. 7**), with an increase in QV (**Supplementary Table 6**). In conclusion, we propose removing all heterozygous variants prior to Merfin as the best practice for polishing pseudo-haploid and haploid assemblies.

In addition, we validated our results using gene annotations, which are sensitive to consensus accuracy error, and particularly to frameshift errors caused by indel errors. We performed *de novo* gene annotation using RefSeq gene annotation pipeline (GNOMON)^37^ on the VGP assemblies polished with the conventional VGP pipeline (v1.6) and compared against assemblies where Merfin was applied at every polishing step. In GNOMON, if a protein alignment supports a predicted model with an indel introducing frameshift or premature stop codons, the model gets labeled as ‘low quality’ with the frame marked for correction. If more than 1 in 10 coding genes in an assembly require corrections, the assembly is excluded from RefSeq. Almost all rejected assemblies used ONT or PacBio CLR reads as their backbone, based on sequence technology information provided by the submitters.

Again, Merfin substantially reduced the number of genes affected by frameshifts, cross-validating the QV and QV* results (**Fig. 4d-f**, **Supplementary Table 7** and example in **Supplementary Fig. 8**). In particular, premature stop codons were significantly reduced with respect to the default polishing in all cases (**Fig. 4d**), with 42.9%, 42% and 21.7% reduction in fArcCen1, rGopEvg1 and bTaeGut1, respectively. Ultimately, 1% or less of genes had code breaks in all cases when using Merfin. Frameshifts were also positively affected (**Fig. 4e**), with 38%, 49.6% and 19.5% reductions in fArcCen1, rGopEvg1 and bTaeGut1, respectively. Less than 3% of genes had frameshifts in all cases when using Merfin. Similarly, the number of protein-coding gene predictions labelled as ‘low quality’ were reduced (**Fig. 4f**). From these results, we now include Merfin as part of the VGP pipeline (v1.7) for its beneficial effect.

Consistent with the variant filtering for genotyping, the improvements in QV with Merfin superseded any hard-filtering attempt using variant call quality score (QUAL) cutoffs at the Arrow polishing step (**Fig. 5a-c, Supplementary Table 8**). For the primary assembly, QV* estimates were consistently higher than the best results attainable by hard-filtering (fArcCen1: Q32.5 vs. Q31.9 at QUAL≥18, rGopEvg1: Q38.7 vs. Q36.7 at QUAL≥21, bTaeGut1: Q44.4 vs. Q42.4 at QUAL≥21). The best QUAL cutoff was not necessarily consistent between species, indicating that a single cutoff cannot produce the best outcome in all cases. The alternate assembly (i.e. alternate haplotype) behaved similarly to the primary assembly, again with Merfin always performing best (fArcCen1: Q31.6 vs Q31.1 at QUAL≥23, rGopEvg1: Q35.2 vs. Q34.2 at QUAL≥26, bTaeGut1: Q42.0 vs. Q40.6 at QUAL≥23); however, it notably differed in best QUAL cutoff values to maximize QV. At increased QUAL cutoffs, both genuine and erroneous corrections are filtered out. Thus, hard-filtering cutoffs perform best when the number of errors corrected exceeds the number of errors introduced at maximum. In contrast, variants selected b Merfin had a wide range of quality scores, with the majority containing higher quality scores yet including some below 25 (**Fig. 5d-f**). Notably, a significant fraction of variants with the highest qualit score assigned were introducing error *k*-mers and thus were rejected by Merfin. Potentially, accumulated sequencing biases in long-reads could lead to erroneous variant calls but can be filtered with more accurate k-mers from short-reads. No hard-filtering methods were able to achieve QV improvements in polishing as observed with Merfin.

**Fig. 5.**
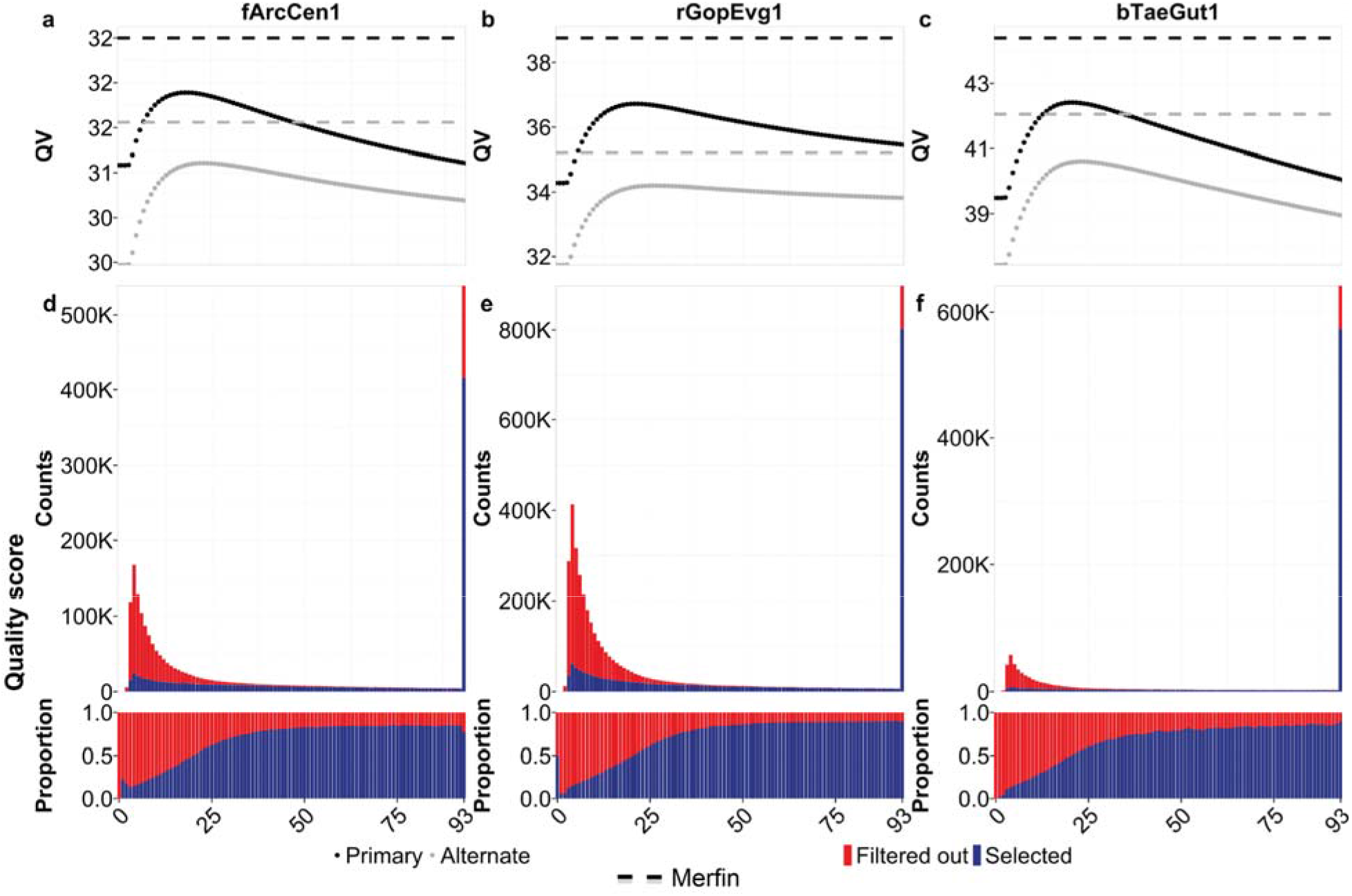
Merfin results against quality scores. **a-c**, QV after polishing as a function of hard-filtered quality scor cutoff in primary (black) and alternate (gray) assembly. Results achieved with Merfin are represented by t horizontal lines for comparison. **d-f**, Number and proportion of variants by quality score selected by Merfin.

## Discussion

Here, we described and demonstrated Merfin, a *k*-mer based tool to evaluate and filter variant calls for improved genotyping accuracy and polishing. Importantly, Merfin allows an innovative alignment-free evaluation and filtering of variants from any VCF generated from any dataset or variant calling method. Merfin always successfully removes false positive calls, superseding any hard-filter based cutoff for both genotyping and polishing. Contrary to the *plateau* effect usually observed in traditional polishing, our *k-* mer-informed polishing is essentially a monotonic function, predicted to improve the consensus accuracy until no more useful variants are produced by the variant caller. This lets polishing pipelines have a natural stopping condition to set, i.e. to stop iterative polishing when less than a certain fraction of produced variants are being filtered.

In addition, using the copy number agreement between the reads and the assembly, we revised *K^*^*, a metric to identify and analyze local expansions and collapses at each *k*-mer genome-wide. We devised QV* and *K^*^* completeness, new quality metrics that account for over and under represented *k*-mers undetected by previous methods^16^. We demonstrated on a complete human genome that our approach allows orthogonal validation of both consensus sequence and variants with any sequencing read data type.

Similarly to all *k*-mer-based estimates, *K^*^* estimates are influenced by the choice of *k*, which is dependent on the quality of the reads. The results presented here assume high-accuracy reads for evaluation and variant filtering, and may therefore not work best with k-mers derived from noisy long-reads (i.e. CLR reads and early ONT data). Presence of sequencing biases will also result in biased *K^*^*, such as the GC bias in Illumina reads or the GA dropouts in HiFi reads^25^. Yet, these effects are limited only to certain regions of the genome, and it could be potentially further mitigated by methods that correct sequencing reads for known biases^38^. Obviously, the ultimate solution will come with less biased, or even unbiased, sequencing reads.

In parallel, the completeness of the assembly also affects the *K^*^*. Pseudo-haploid or haploid representation of a genome may potentially lead to suboptimal evaluation because of the missing copies. However, we argue this is a limitation of the assemblies themselves, rather than a limitation of the methods used to evaluate and polish them. While haploid or partially phased (e.g. FALCON-Unzip^36^) assemblies can be preferred for some applications, a faithful reconstruction of the complete genome should be preferred for both evaluation and comparative purposes, as well as for many biological analyses that can benefit from the presence of both haplotypes (e.g. trio binning^33,39^). Representing a diploid genome as a haploid or pseudo-haploid assembly introduces complications in the evaluation, since the *k*-mers in the consensus will not fully reflect the *k*-mers in the read set. Homozygous *k*-mers will be underrepresented, and some of the alternate haplotype *k*-mers will be completely missing. The recent developments in assembly graphs enable the representation of complete haplotypes with enhanced accuracy and completeness^40^, suggesting that assembly tools and state-of-the-art assemblies are moving toward this direction. If this condition is met, the information contained in the reads can be fully harnessed to evaluate and improve genome assemblies.

Merfin presents the first *k*-mer based variant filtering to the best of our knowledge, enabling higher precision in genotyping and improving assembly accuracy. This will become critical particularly in medical genomics and many other applications, where reliable genotyping is highly important. Polishing with Merfin will rescue assemblies built from noisy long reads, when the more recent, accurate platforms are not accessible, or when sequencing biases are subject to correction using complementary sequencing platforms.

## Online methods

### Genotyping benchmark

Variant calls from HG002 submitted to precisionFDA Truth Challenge^1^ were downloaded from https://data.nist.gov/od/id/mds2-2336. In brief, ~35x Illumina PCRfree, ~36x PacBio HiFi, and ~47x ONT reads were aligned to the human genome reference (GRCh38) with no alternates. Variants were called with GATK HaplotypeCaller v4. Unfiltered and hard-filtered set was downloaded and benchmarked with hap.py (v0.3.12-2-g9d128a9) following the best practices from a previous study^26^. Precision and recall were collected before and after filtering the variants with Merfin.

PCR-free Illumina paired-end reads (2×250 bp) were obtained from NIST (https://ftp-trace.ncbi.nlm.nih.gov/giab/ftp/data/AshkenazimTrio/HG002_NA24385_son/NIST_Illumina_2x250bps/) and 21-mers were collected using Meryl. *K*-mers with frequency > 1 were used as read *k*-mers to avoid *k*-mer collisions from sequencing errors and improved computational performance.

### Revised K*

*K*-mers are substrings of length *k* of a given DNA sequence. Given the assembly consensus sequence, we compute all its constituent *k*-mers. Similarly, we compute all *k*-mers represented in a set of WGS reads from the same individual. We then ask how the frequency of each *k*-mer in the read set is mirrored in the assembly *k*-mer set. If the read set is a faithful representation of the genome (i.e. in the absence of random DNA sampling and sequencing biases), then the closer the consensus sequence is to the read set, the closer it is also to the genome the reads were generated from. This principle can be usefully represented by our revised K*, where for each *k*-mer in the consensus we can calculate (**Fig. 1a**):

K_C_ = *k*-mer count in the consensus sequence
K_R_ = *k*-mer count in the read set

To account for the uncertainty associated with the underlying Poisson sampling process, in any sequencing experiment the read set covers on average the original genome multiple times. It is therefore useful to determine the expected copy number of a particular *k*-mer in the assembly given the read set, K_r_, as:

*c* = haploid peak from K_R_ histogram
K_R_ = the *k*-mer count expected in the consensus based on the read set, i.e. ⍰ K_R_ / *c* ⍰

Note that K_r_ - K_C_ expresses the number of copies of any particular *k*-mer that is underrepresented (collapsed; positive value) or overrepresented (expanded; negative value) in the assembly.

With these definitions, we can now define K* as:

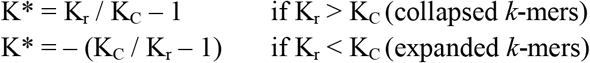

Which can be reduced to:

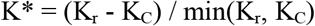

Note that K* converges to 0 if the *k*-mer frequency in the assembly matches the expected copy number in the reads. Missing *k*-mers (i.e. found in the assembly but not in the read set) have a special behavior with K* being “undefined” for K_r_ = 0.

### Probabilistic K-mer copy-number estimation

To estimate *k*-mer copy-number in the genome, we modified Genomescope2^27^ to obtain the associated probability at each K_R_. Our additions were subsequently integrated in the current version of Genomescope2 (https://github.com/tbenavi1/genomescope2.0). Unmodified fitted model 1-to 4-copy *k*-mer distributions were used to infer the probability that a particular *k*-mer frequency observed in the read set implied a particular copy *k*-mer in the genome. Using this model, Merfin provides a script generating a lookup table for each k-mer frequency in the raw data with the most plausible *k*-mer multiplicity and its associated probability (https://github.com/arangrhie/merfin/tree/master/scripts/lookup_table).

### QV estimation using the K*

An average genome-wide QV* is obtained by counting all k-mers not present compared to the expected copy number estimated from the read set. We collect all k-mers excessively found in the assembly (K_E_) and estimate the error rate given all k-mers in the assembly (K_total_).

K_E_ = Σ K_C_ - K_r_ when K_C_ > K_r_ for all positions in the assembly

The Phred-scaled QV* can be computed using the implementation in Merqury^16^ or YAK^28^.

We follow the implementation in Merqury and compute the probability P that a base in the assembly is correct and in its expected frequency:

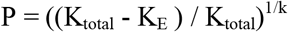

Which leads to error rate E being:

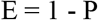

Hence the phred scaled QV* becomes:

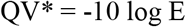

### Assembly completeness using the K*

To estimate completeness, we collect all k-mers that should be present but are absent from the assembly. Unlike Merqury, we account for the k-mer frequency and count any k-mer that should be added to meet the expected frequency from the reads K_A_.

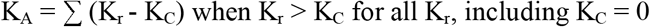

We compute the completeness Comp given all K_r_:

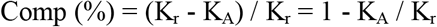

### Sequence data

For the HG002 results, data can be found at https://github.com/human-pangenomics/HG002_Data_Freeze_v1.0. For the VGP datasets, PacBio CLR and 10x Genomics linked reads can be found at https://vgp.github.io/genomeark/^5^.

### Evaluation of CHM13 assemblies

All scripts used for CHM13 evaluation can be found here: https://github.com/gf777/misc/tree/master/merfin. Briefly, we generated genome-wide K* tracks using Merfin option -dump (merfin_dump.sh). *K*-mer counts databases for both the assemblies and the raw Illumina and HiFi reads were computed using Meryl (https://github.com/marbl/meryl). Peak values of 106.8 and 31.8 derived as the kcov value from Genomescope2 were used for Illumina and HiFi *k*-mers, respectively. The tracks were converted to bigWig and loaded in IGV^41^ for visualization. We used a custom script (simplify_dump.sh) to count the number of bases with the same K* values for both Illumina and HiFi *k*-mers, which were then used to generate the genome-wide K* comparison. The titration experiment was performed downsampling the reads with the ‘seqtk sample’ command (https://github.com/lh3/seqtk).

### Variant calling and polishing of HG002 assemblies

Variant calling and polishing of HG002 assemblies was performed using medaka 1.2.6 (https://github.com/nanoporetech/medaka) using the models specified in **Supplementary Table 3** for each dataset. Medaka was first run in the consensus mode (medaka_consensus) and subsequently in the variant mode (medaka_variant) to generate the vcf of the variant calls. Medaka filtered variant set was then used in conjunction with bcftools v1.9 in the consensus mode with the -H 1 option to generate a consensus sequence. The same procedure was followed for the Merfin assemblies, except that Merfin was used to filter Medaka vcf prior to consensus generation. Polishing was repeated twice, and in the second round the assembly polished with Merfin was used as reference.

### Assembly-based small variant calling assessment

We used dipcall (https://github.com/lh3/dipcall) to generate the small variants from the assembly. Dipcall takes a diploid assembly and a reference genome to produce a variant call file (VCF) that contains all variants that are present in the assembly compared to the reference. We then compared the variant calls against GIAB truth set v4.2.1 using hap.py (https://github.com/Illumina/hap.py). The GIAB variant calling truth set for HG002 sample can be found in: (ftp://ftp-trace.ncbi.nlm.nih.gov/giab/ftp/release/AshkenazimTrio/HG002_NA24385_son/NISTv4.2.1/). We used the following commands for the evaluation:

~~~
./run-dipcall <output_prefix> <GRCh38.fa> <pat.fa> <mat.fa> -t 8 -x hs38.PAR.bed
docker run -it pkrusche/hap.py:latest /opt/hap.py/bin/hap.py HG002_GRCh38_1_22_v4.2.1_benchmark.vcf.gz DIPCALL_OUTPUT.dip.vcf.gz -f HG002_GRCh38_1_22_v4.2.1_benchmark_noinconsistent.chr20.bed -r GCA_000001405.15_GRCh38_no_alt_analysis_set.fna -o OUTPUT --pass-only --engine=vcfeval -- threads=32
~~~

### Variant calling and polishing of VGP assemblies

While the original assemblies were generated with different versions of the VGP pipeline^5^ (https://github.com/VGP/vgp-assembly/tree/master/pipeline), to polish the assemblies of the flier cichlid (v1.0), the Goode’s thornscrub tortoise (v1.5), and the zebra finch (v1.6) with Merfin we used the VGP pipeline v1.6 (**Supplementary Fig. 7**). In the first round of polishing, long Pacbio CLR reads were aligned with pbmm2 v1.0.0, variants were called with variantCaller v2.3.3 (arrow) with the ‘-o ${asm}.vcf’ option. A custom script (https://github.com/arangrhie/merfin/blob/master/scripts/reformat_arrow/) included in Merfin was used to properly format the vcf file. 21-mer databases for both the assemblies and the 10x linked-reads were generated with Meryl. 10x barcodes were trimmed from the reads using the script available in Meryl. The haploid 21-mer coverage and the lookup tables were computed using our modified Genomescope2 script included in Merfin:

~~~
Rscript $merfin/lookup.R ${asm}.21.meryl.hist 21 ${asm}.21.lookup 2
~~~

Similarly to HG002, the consensus was generated with bcftools v1.9 using the filtered vcf generated by Merfin. The same strategy was applied for the other polishing steps, except that Longranger v2.2.2 was used for mapping the 10x Genomics linked-reads and Freebayes v1.3.1 for variant calling. For Variant calling and polishing of zebra finch trio, Freebayes calls were filtered using Bcftools v1.9 with the - i’(GT=“AA” || GT=“Aa”)’ option prior to Merfin filtering. *K*-mer counts databases for both the assemblies and the raw Illumina reads were computed using Meryl, and Merfin was run with a peak value of 35.2 derived as the kcov value from Genomescope2.

### Evaluation of the assemblies

QV and phasing analyses of HG002 and zebra finch trios were performed using Merqury^16^ (https://github.com/marbl/merqury/) in the trio mode using 21-mers and default parameters. Similarly, primary and alternate scaffolds of the VGP assemblies were separated and Merqury QV was estimated on both using 21-mers and default parameters.

### Gene annotation of VGP assemblies

Annotation was performed as previously described^5^, using the same transcript, protein and RNA-Seq input evidence for the annotation of the unpolished, polished and Merfin assemblies of each species. For *Taeniopygia guttata*, a total of 100,000 *Taeniopygia guttata* ESTs, GenBank and known RefSeq and 10 billion same-species reads for over 13 tissues were aligned to the genome, in addition to all Genbank *Aves* proteins, known *Aves*, human and *Xenopus* RefSeq proteins, and RefSeq model proteins for *Parus major*, *Gallus gallus*, *Columbia livia* and *Pseudopodoces humilis*. For *Gopherus evgoodei*, 1.22 billions RNA-Seq reads from 5 tissue types from *Gopherus* and *Chelonoidis* species were aligned to the assemblies in addition to all known RefSeq proteins from human, *Xenopus,* and *Sauropsida*, and model RefSeq proteins from *Chrysemys picta*, *Pelodiscus sinensis*. For *Archocentrus centrarchus*, 476 million same species RNA-Seq reads from 9 tissue types were aligned to the assemblies in addition to all *Actinoipterygii* GenBank proteins, human and *Actinopterygii* known RefSeq proteins and *Oryzias latipes*, *Oreochromis niloticus*, *Monopterus albus*, *Xiphophorus maculatus* model RefSeq proteins.

#### Plots

A combination of bash and Rscript was used for data analysis and visualization. All scripts used to generate the plots are available at https://github.com/gf777/misc/tree/master/merfin/paper/figures.

## Availability

A stable release and the C++ source code for Merfin, and examples from this work are available under Apache License 2.0 (https://github.com/arangrhie/merfin). The only dependency is the *k*-mer counter Meryl, which comes with the release. Merfin can be run in five modes: 1) the -filter mode scores each variant, or variants within distance *k* and their combinations by error k-mers for improved genotyping; 2) the -completeness mode generates completeness metrics; 3) the -dump mode computes K_C_, K_R_, K* for each base in the assembly along with QV and QV* for each sequence; 4) the -hist mode provides a K* histogram and genome-wide QV and QV* averages; 5) the -polish mode scores each variant, or variants within distance *k* and their combinations by the K* for polishing. Merfin is fully parallelized using OpenMP. The K* tracks obtained from each platform for the CHM13 v1.0 assembly are available in the associated UCSC browser (http://t2t.gi.ucsc.edu/chm13/dev/hub.txt).

## Author Contributions

A. R., G. F., B. P. W. and E. M. conceived and implemented Merfin. G. F. and A. R. performed the validation analyses. K. S. performed the GIAB variant calling analysis on HG002. F. T. generated the gene annotations for the VGP genomes. S. K. contributed to the conceptual development. G. F. and A. R. wrote the manuscript. G. F., A. R., E. D. J. and A. M. P. conceived the study. All authors reviewed, edited, and approved the manuscript.

## Competing Interests statement

The authors declare no competing interest.

## Acknowledgments

We thank T. Rhyker Ranallo-Benavidez and Michael C. Schatz for the useful discussion on adapting Genomescope2 models. We also thank the communities of the T2T, HPRC and VGP consortia for their constant support. G. F. and E. D. J. were supported by Rockefeller University and HHMI funds. A. R., B. P. W., S. K, and A. M. P. were supported by the Intramural Research Program of the National Human Genome Research Institute, National Institutes of Health. The work of F.T-N was supported by the Intramural Research Program of the National Library of Medicine, National Institutes of Health. K. S. was supported by NIH/NHGRI (R01HG010485, U41HG010972, U01HG010961, U24HG011853, OT2OD026682). E. W. M. was partially supported by the German Federal Ministry of Education and Research (01IS18026C)

## Supplementary Figures

**Supplementary Figure 1.**
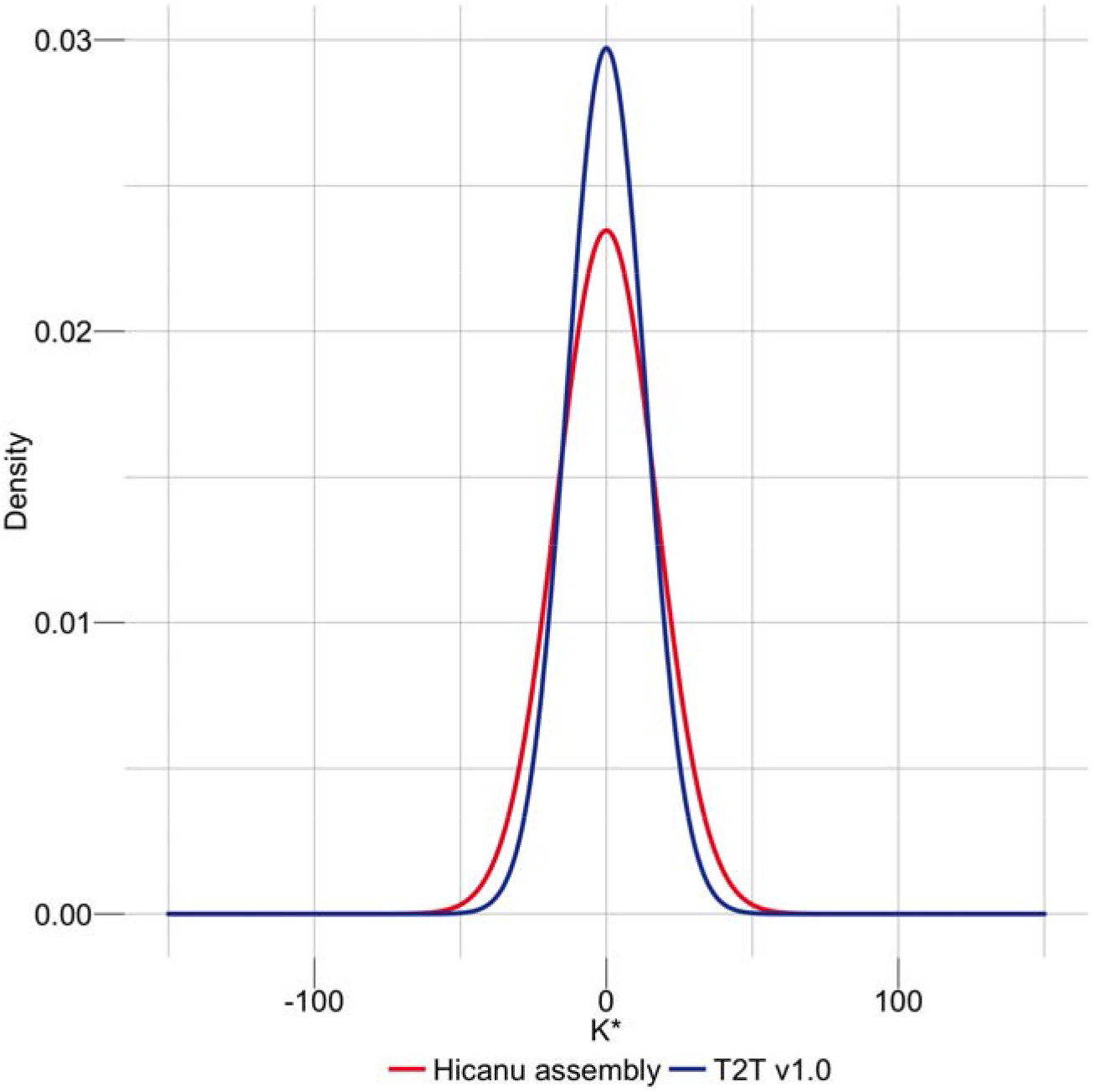
Genome-wide density distribution of the K* using Illumina*k*-mers. When the assembly is in agreement with the raw data, the K* is normally distributed with mean 0, and th smaller the standard deviation the higher the agreement. CHM13 v1.0 shows a less dispersed distribution of the K* compared to a regular HiCanu assembly.

**Supplementary Figure 2.**
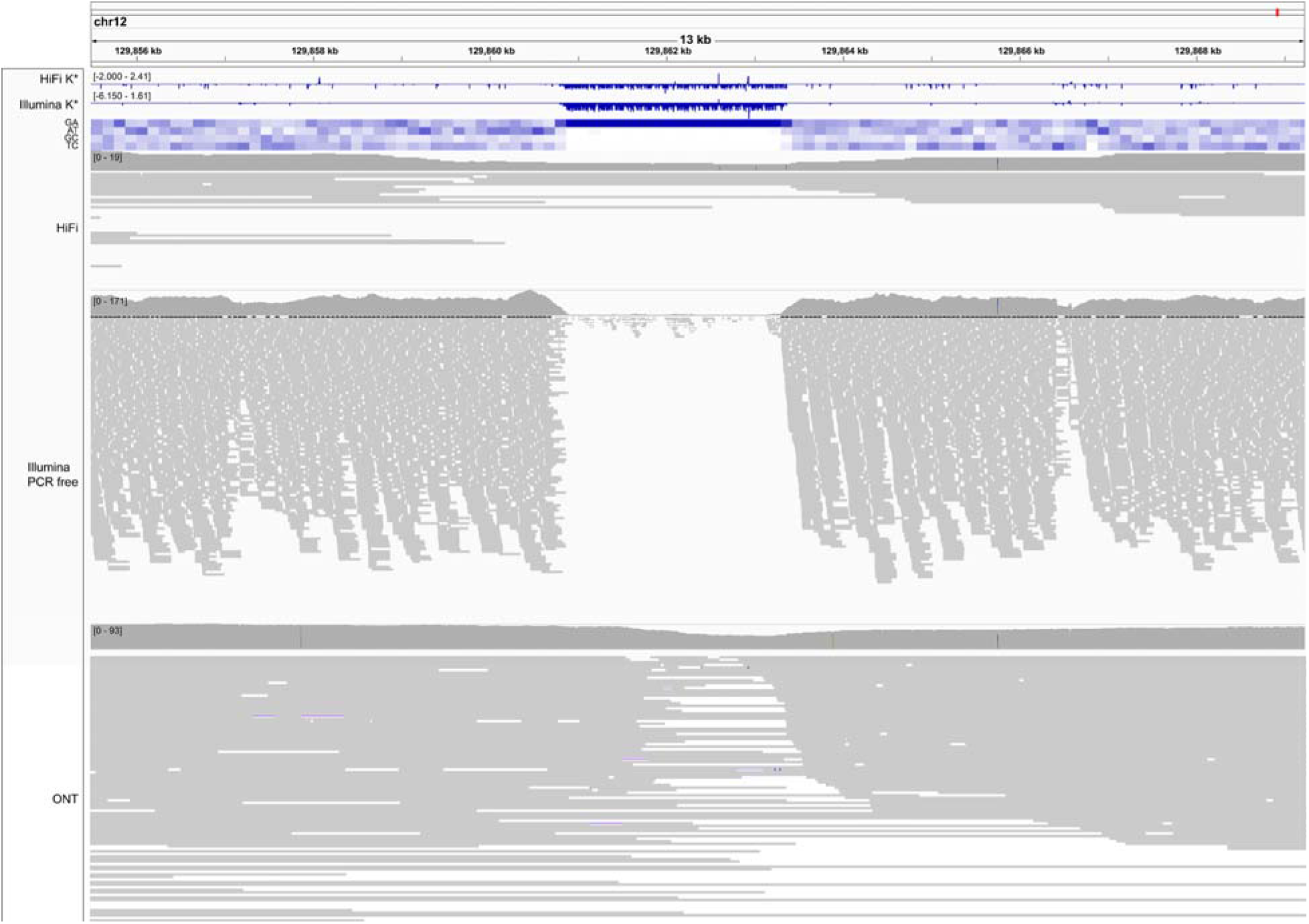
A region of negative K* highlighting sequencing bias. An example of low coverage in both HiFi and Illumina reads associated with high guanine content, and specifically a GA-rich repeat (heatmap). GA bias has been reported in Pacbio HiFi data, and results in gaps in the assembly that in CHM13 were filled with Nanopore data^22^. The K* both from HiFi and Illumina *k*-mers (top tracks) recapitulate the coverage drop. Nanopore coverage appears less affected. Position Chr12:~129,862,000 bp.

**Supplementary Figure 3.**
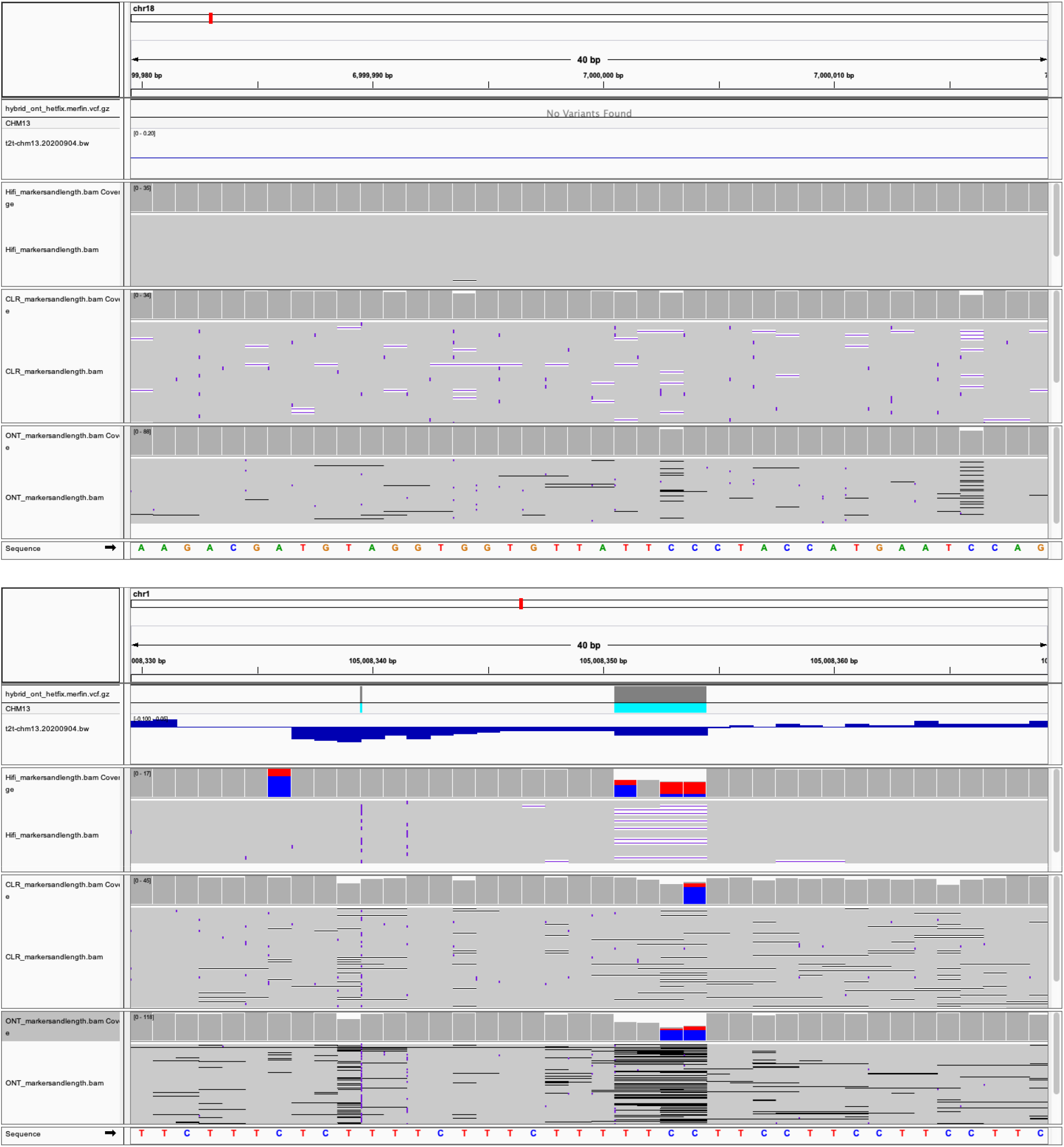
The K* can identify issues in the assembly at the base level. **a**, 40 bp window with K* close to 0, highlighting perfect agreement of the assembly with the raw reads. Position Chr18:~7,000,000 bp. **b**, A region of negative K* in coincidence with two heterozygous indels. Position Chr1:~105,008,350 bp.

**Supplementary Figure 4.**
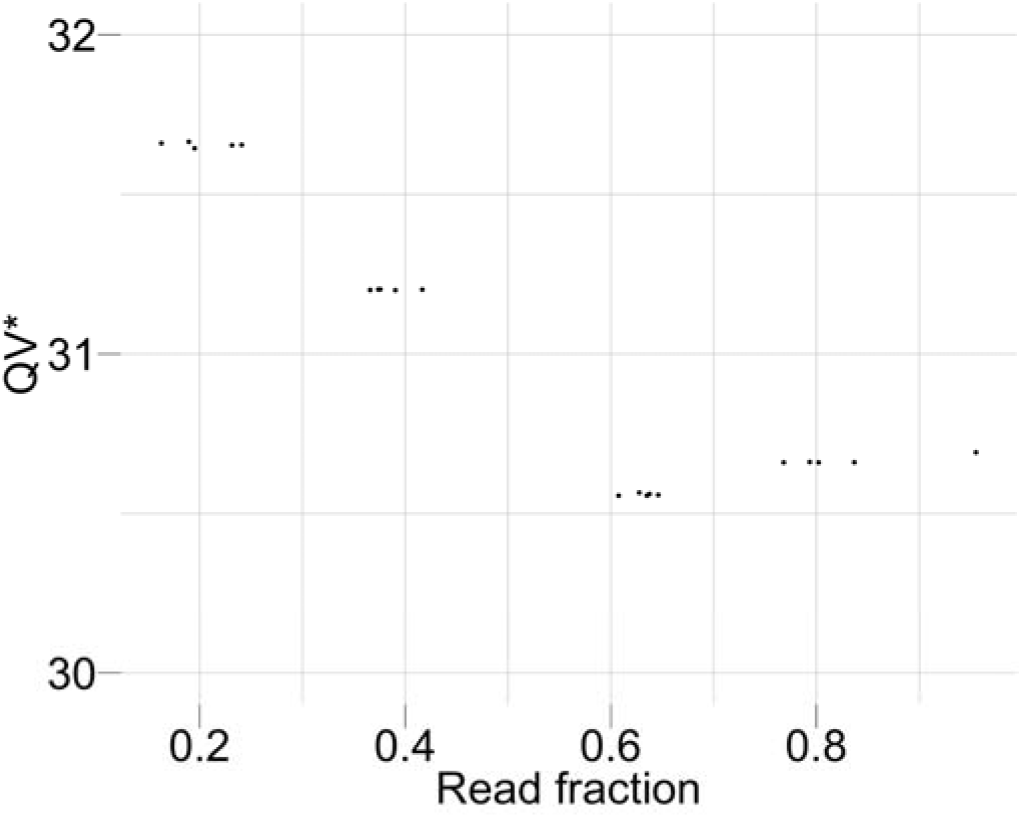
Coverage titration experiment and impact on QV*. The QV* is only marginally influenced by the coverage of the dataset being considered.

**Supplementary Figure 5.**
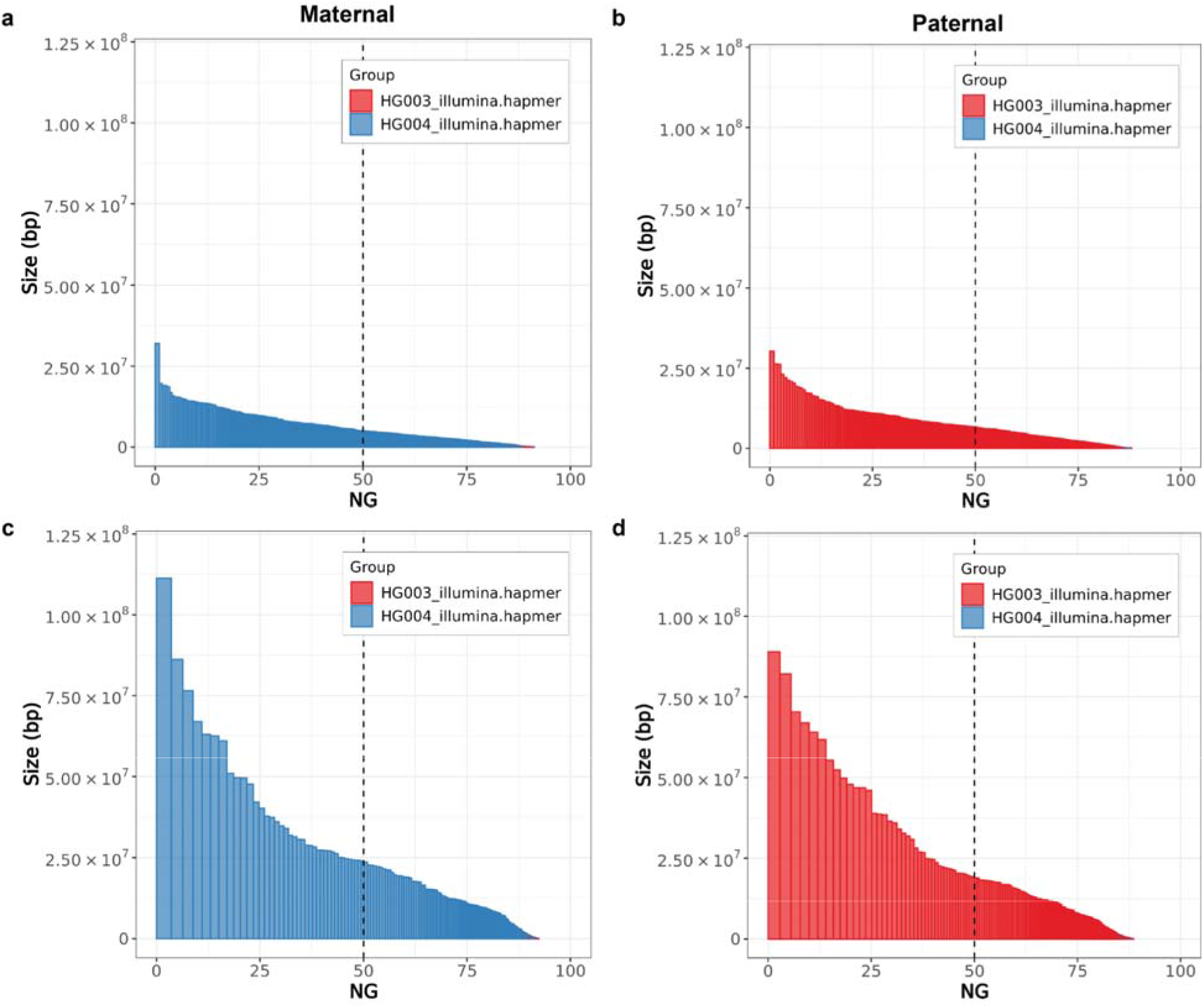
Haplotype phasing before and after polishing with Merfin. In both parental assemblies, the haplotypes remained fully phased, and the size of the blocks significantly increased compared to the unpolished version (**a,b**) after polishing with Merfin (**c,d**). A theoretical human genome size of 3.1 Gbp was used to normalize NG* values.

**Supplementary Figure 6.**
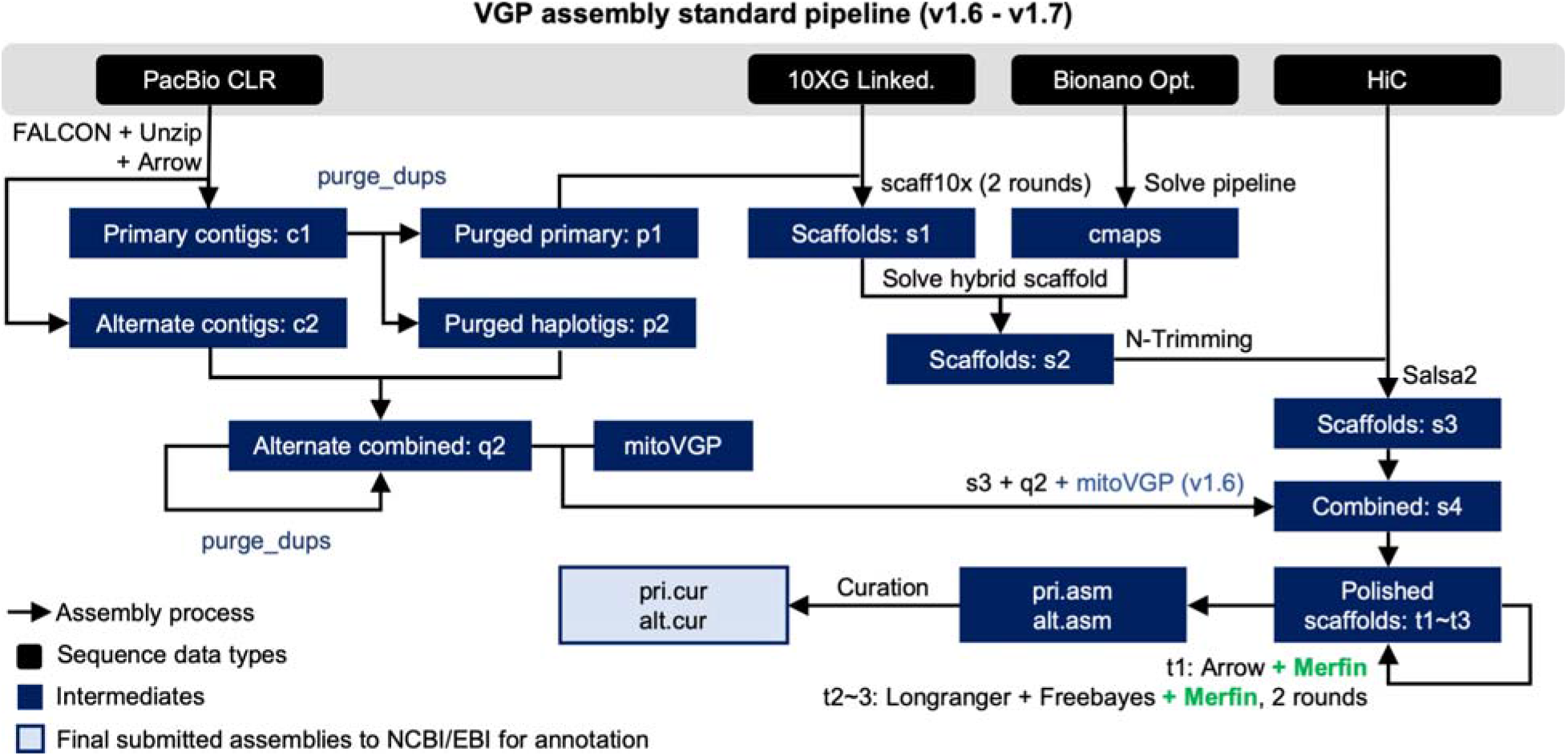
VGP assembly pipeline. Compared to the previous v1.6, the introduction of Merfin in v1.7 (green) resulted in a minimal change of the workflow, but in a generalized improvement in QV scores and gene annotations. Pipeline available at https://github.com/VGP/vgp-assembly.

**Supplementary Figure 7.**
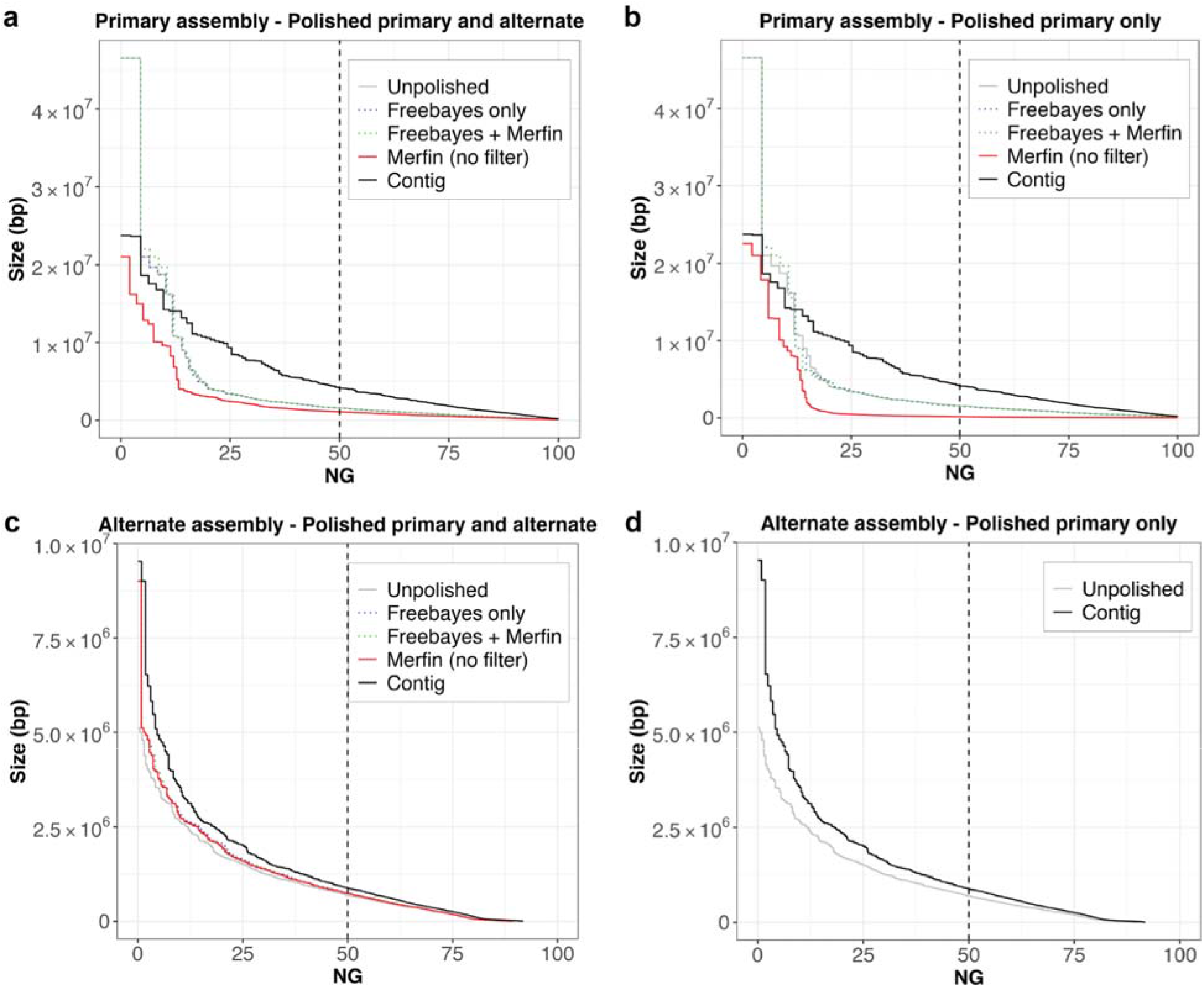
Phase block analysis of zebra finch pseudo-haplotype assembly. **a**, Phase blocks in the primary assembly after mapping the reads to both the primary and alternate assemblies. **b,** Phase blocks in the primary assembly after mapping the reads to both the primary only. **c**, Phase blocks in the alternate assembly after mapping the reads to both the primary and alternate assemblies. **d**, Phas blocks in the alternate assembly. In all cases, the application of Merfin filtering minor heterozygous variants (green) leads to block sizes better or comparable to prior polishing methods alone (blue). Unpolished assembly in gray. Results of Merfin without filtering in red. A genome size of ~1.03 Gbp derived from Genomescope2 was used to normalize NG* values.

**Supplementary Figure 8.**
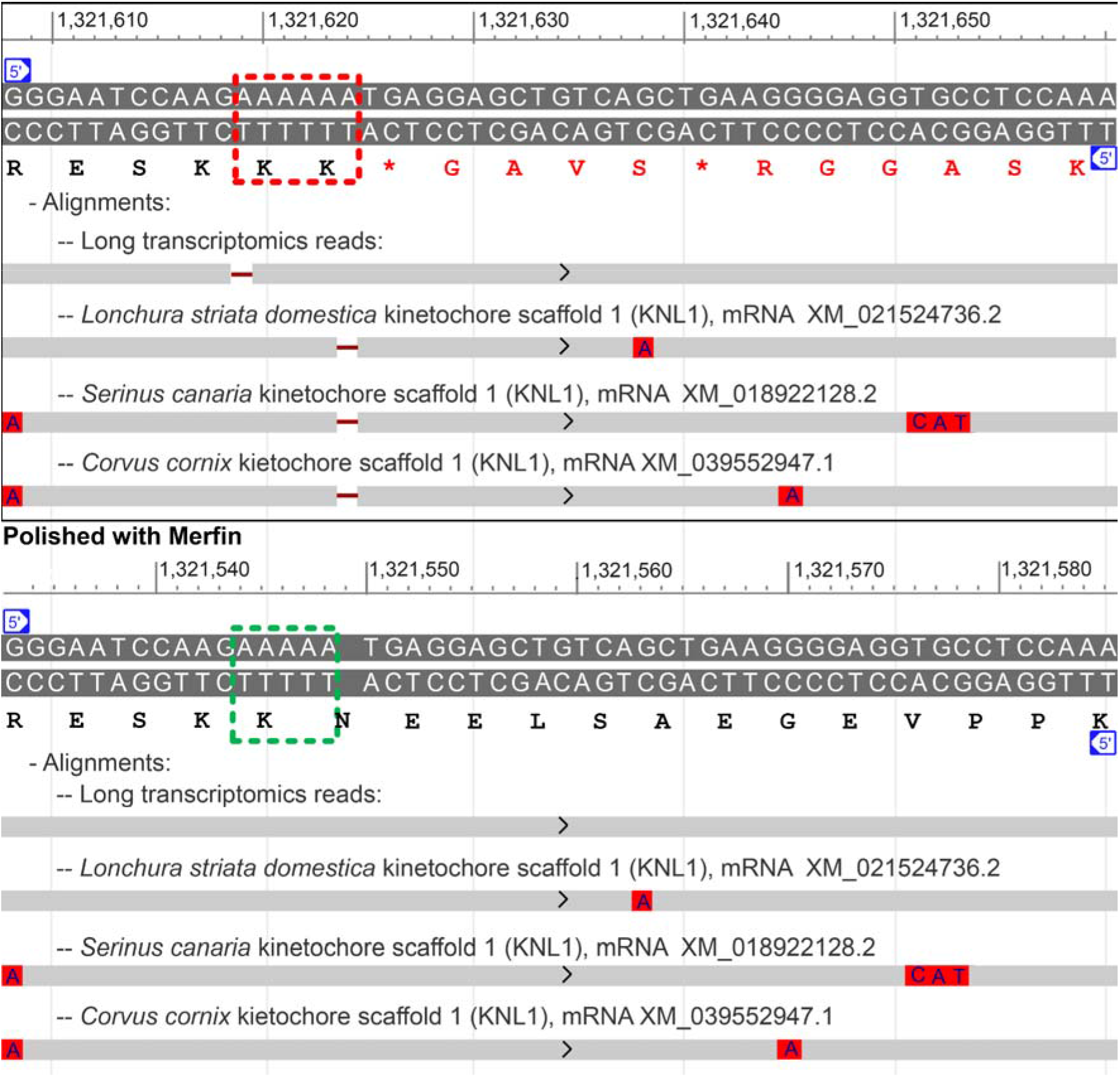
Effect of merfin correction on the kinetochore scaffold 1 protein (KNL1) annotation. **a**, Deleterious presence of an extra A around position 1,321,620 of scaffold_7 (red box) in the polished, non-merfin-corrected sequence is indicated by a 1-base gap in the alignments of zebra finch PacBio IsoSeq SRR8695295.20794.1 and KNL1 transcripts from three other Passeriformes songbirds. This insertion causes a disruption in the frame and a premature stop codon in the translated sequence (se amino acid sequence in red). **b**, Corresponding span in the merfin-corrected assembly, with gapless alignments of the IsoSeq read and Passeriformes transcripts, and uninterrupted translation.

## Notes

https://github.com/arangrhie/merfin

### Competing Interest Statement

The authors have declared no competing interest.

